# Effects of Glutamate Delta 1 Receptor (GluD1) Deletion on the Ultrastructural Features of Corticostriatal and Thalamostriatal Synapses in Mice

**DOI:** 10.64898/2026.05.26.727726

**Authors:** Larissa Yue, Karina Dalal, Shashank M. Dravid, Yoland Smith, Rosa M. Villalba

**Affiliations:** Emory College of Arts and Sciences, Emory University, Atlanta, Georgia, USA; Morehouse School of Medicine, Atlanta, Georgia, USA; Department of Psychiatry and Behavioral Science, Texas A&M University, College Station, Texas, USA; Department of Neurology, Emory University, Atlanta, Georgia, USA; Emory National Primate Research Center, Emory University, Atlanta, Georgia, USA

**Keywords:** electron microscopy, vGluT1, vGluT2, striatum, receptor, synaptic plasticity

## Abstract

The glutamate delta 1 receptor (GluD1) represents a unique subtype of ionotropic glutamate receptors that is strongly expressed in the mammalian striatum. Disruptions of the *GRID1* gene, which encodes GluD1, have been associated with neuropsychiatric disorders, including schizophrenia and autism spectrum disorder; however, the role of GluD1 in the brain remains poorly understood. Previous studies in mice have demonstrated that the knockout of striatal GluD1 led to fear-conditioning deficits and depressive-like behaviors. Furthermore, these mice exhibited reduced excitatory input to the striatum due to a loss of thalamostriatal innervation, whereas corticostriatal innervation was unaffected. In this study, we examined whether changes in synapse morphology contribute to the observed functional deficits. We found that the ablation of GluD1 does not affect synaptic targeting patterns of corticostriatal and thalamostriatal terminals, using transmission electron microscopy. We further utilized three-dimensional reconstruction to obtain quantitative data on synapse ultrastructure and found no significant changes in corticostriatal and thalamostriatal synaptic components, including the presynaptic terminal volume, postsynaptic density area and morphology, and postsynaptic dendritic spine volume. These findings support a model in which GluD1 regulates input-specific circuit organization and synaptic connectivity rather than the structural morphology of individual synapses.

## 1 INTRODUCTION

Glutamate is the principal excitatory neurotransmitter in the central nervous system and regulates a wide range of functions, including cognition, memory, and learning. Its effects are mediated through two main classes of receptors: metabotropic (mGluRs), which modulate intracellular signaling through G-proteins, and ionotropic receptors, which function as ligand-gated ion channels. The ionotropic family includes the well-characterized AMPA, NMDA, and kainate receptors, as well as the glutamate delta receptors (GluD1 and GluD2). Although GluD receptors are structurally related to other ionotropic glutamate receptors (Wo & Oswald, 1995), they do not bind glutamate nor exhibit conventional ligand-gated ion channel activity (Kakegawa et al., 2011; Kristensen et al., 2016; Lomeli et al., 1993). Instead, accumulating evidence indicates that GluD receptors play critical roles in synapse formation and maintenance (Choi et al., 2026; Matsuda et al., 2010; Uemura et al., 2010).

Specifically, GluD1 is highly enriched in brain areas implicated in the pathophysiology of neuropsychiatric disorders (Choi et al., 2025; Conley et al., 2025; Gandhi et al., 2021; Gantz et al., 2020; Khamma et al., 2022; Liu et al., 2020). Consistent with this distribution, genetic studies have identified strong associations between disruptions of the *GRID1* gene, which encodes GluD1, and several psychiatric conditions including schizophrenia (Greenwood et al., 2011; Guo et al., 2007; Treutlein et al., 2009), autism spectrum disorder (Glessner et al., 2009; Griswold et al., 2012; Nord et al., 2011), and major depressive disorder (Edwards et al., 2012). In addition, altered GluD1 expression—driven by mutations in the MECP2 gene—disrupts the critical balance of synaptic development in both Rett syndrome patient-derived neurons and mouse models (Livide et al., 2015; Patriarchi et al., 2016). Deficits in fear conditioning, along with abnormal emotional, social, and depressive-like behaviors were also observed in mice with GluD1 deletion (Benamer et al., 2018; Nakamoto, Kawamura, et al., 2020; Yadav et al., 2012; Yadav et al., 2013). Beyond neuropsychiatric disease, GluD1 has also been implicated in neurological conditions such as chronic pain, addiction, and motor dysfunction (Gandhi et al., 2021; Liu et al., 2018; Liu et al., 2020; Sabnis et al., 2024; Sabnis et al., 2025; Shelkar et al., 2025; Yadav et al., 2012; Yadav et al., 2013). Together, these findings highlight the broad clinical relevance of GluD1 and underscore the importance of understanding its role in neural circuit function.

Lacking conventional ligand-gated ion channel activity, GluD1 instead participates in trans-synaptic adhesion complexes by interacting with presynaptic neurexins (Nxn) proteins via cerebellin (Cbln) released from presynaptic terminals (Kakegawa et al., 2011; Kristensen et al., 2016; Lomeli et al., 1993), forming a GluD–Cbln–Nxn complex that functions as a synaptic organizer (Cheng et al., 2016; Elegheert et al., 2016; Matsuda et al., 2010; Uemura et al., 2010). While this mechanism has been well characterized for its family member GluD2 in the cerebellum (Berridge et al., 2018; Hirano, 2012; Ichikawa et al., 2016; Kashiwabuchi et al., 1995; Nakamoto, Konno, et al., 2020; Pernice et al., 2019; Yuzaki, 2011), the precise role of GluD1 in the mammalian brain remains incompletely understood.

At the anatomical level, previous morphological and ultrastructural studies have shown that GluD1 is highly expressed in the dorsal striatum, where it preferentially localizes at asymmetric axo-dendritic synapses formed onto medium spiny neurons (MSNs) (Hoover et al., 2020; Liu et al., 2020). In this region, excitatory inputs from the thalamus and cortex can be distinguished by their expression of vesicular glutamate transporter type 2 (vGluT2) and vesicular glutamate transporter type 1 (vGluT1), respectively (Fremeau et al., 2001; Fujiyama et al., 2006; Lacey et al., 2005; Raju et al., 2006). Notably, cerebellin-1 (Cbln1), a key component of the tripartite complex, is highly expressed in neurons of the parafascicular nucleus (Pf) of the thalamus (Kusnoor et al., 2010; Miura et al., 2006; Otsuka et al., 2016), which constitute the principle source of thalamic glutamatergic axo-dendritic synapses onto striatal projection neurons (Dube et al., 1988; Raju et al., 2006; Sadikot et al., 1992). This anatomical and molecular organization suggests that GluD1 may play a selective role in regulating thalamostriatal vGluT2 synaptic connectivity.

Consistent with this idea, Liu et al. (2020) demonstrated that conditional deletion of GluD1 in the mouse dorsal striatum, a model designed to selectively eliminate the GluD1 receptor from striatal circuits, results in a significant reduction in excitatory synaptic transmission in MSNs and impaired behavioral flexibility, accompanied specifically by a selective decrease in thalamostriatal vGluT2-immunoreactive (IR) input, while corticostriatal vGluT1-IR input remains unchanged. These findings indicate that GluD1 plays a critical role in thalamostriatal physiology; however, the structural alterations underlying this functional deficit remain unclear. To gain deeper insight into potential structural alterations, we used a GluD1-knockout (KO) mouse model and transmission electron microscopy (TEM) to assess possible modifications of postsynaptic targets of corticostriatal and thalamostriatal terminals. Serial images obtained from high-resolution array tomography/scanning electron microscopy (AT/SEM) and three-dimensional (3D) reconstruction were used to further characterize the ultrastructural features of striatal glutamatergic terminals and their synapses in wild-type (WT) and KO mice. Because synaptic ultrastructure is closely tied to neurotransmitter release, synaptic efficacy, and plasticity (Harris et al., 1992; Harris & Kater, 1994; Holler et al., 2021; Pan et al., 2023; Villalba & Smith, 2010), examining these features provides a means to determine whether reduced excitatory transmission arises from a loss of synapses or from alterations in the ultrastructure of individual synapses.

## 2 MATERIALS AND METHODS

### 2.1 Animals

Three WT mice and three KO mice from Creighton University School of Medicine were used in this study (Table 1). All housing, feeding, and experimental procedures were conducted in accordance with the guidelines for the care and use of laboratory animals established by the National Institutes of Health (National Research Council) and were approved by Creighton University’s Institutional Animal Care and Use Committees (IACUC).

**Table 1:**
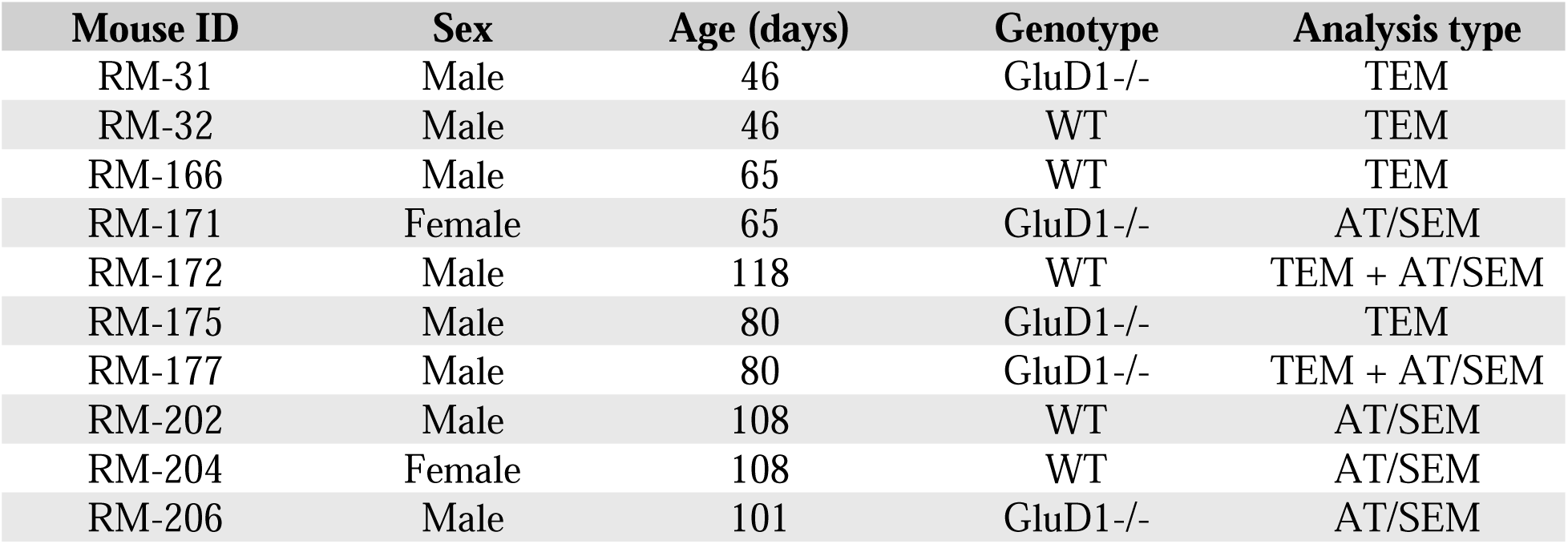
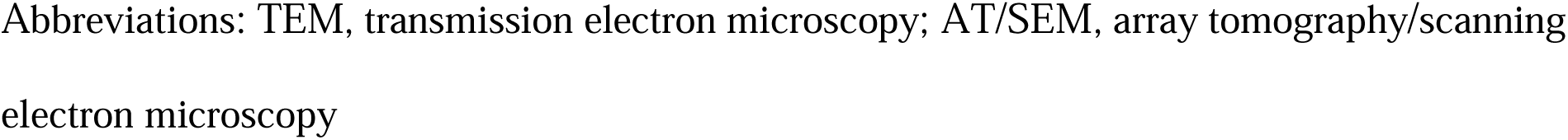
Animals used.

| Mouse ID | Sex | Age (days) | Genotype | Analysis type |
| --- | --- | --- | --- | --- |
| RM-31 | Male | 46 | GluD1-/- | TEM |
| RM-32 | Male | 46 | WT | TEM |
| RM-166 | Male | 65 | WT | TEM |
| RM-171 | Female | 65 | GluD1-/- | AT/SEM |
| RM-172 | Male | 118 | WT | TEM + AT/SEM |
| RM-175 | Male | 80 | GluD1-/- | TEM |
| RM-177 | Male | 80 | GluD1-/- | TEM + AT/SEM |
| RM-202 | Male | 108 | WT | AT/SEM |
| RM-204 | Female | 108 | WT | AT/SEM |
| RM-206 | Male | 101 | GluD1-/- | AT/SEM |

Abbreviations: TEM, transmission electron microscopy; AT/SEM, array tomography/scanning electron microscopy

At the time of euthanasia, mice were deeply anesthetized with isoflurane and transcardially perfused with Ringer’s solution, followed by a fixative solution containing 4% paraformaldehyde and 0.1% glutaraldehyde. After perfusion, brains were removed from the skull and post-fixed in 4% paraformaldehyde. Tissue was then sectioned into 60 µm-thick coronal sections using a vibrating microtome and stored at -20℃ in an anti-freeze solution until immunohistochemical processing.

### 2.2 Antibodies

All primary antibodies used in this study are commercially available, well characterized, and registered in the Research Resources Identifiers (RRIDs) Portal (Table 2). The specificity of the GluD1 antibody has been further validated by the absence of immunostaining in the striatum of KO mice (Liu et al., 2020).

**Table 2:** List of primary antibodies used.

| Antibody | Cat. Number | Host | Vendor | Dilution | RRIDs |
| --- | --- | --- | --- | --- | --- |
| vGluT1 | Ab5905 | Guinea pig | Millipore | 1:5,000 | AB_2301751 |
| vGluT2 | VGT2-3 | Rabbit | Mab Technologies | 1:5,000 | AB_2315568 |

### 2.3 vGluT1 and vGluT2 immunocytochemistry and tissue preparation

#### 2.3.1 Electron microscopy pre-embedding immunoperoxidase

Striatal vibratome sections from 3 WT and KO mice were processed to identify vGluT1 and vGluT2-IR terminals at the electron microscopic level. Sections were first incubated in a sodium borohydride solution (1% in PBS; 0.01 M, pH 7.4) at room temperature (RT) for 20 min, followed by five washes in PBS. Sections were then submerged in a cryoprotectant solution (25% sucrose and 10% glycerol in PB; 0.05 M, pH 7.4) at RT for 20 min before frozen at -80 ℃ for 20 min, thawed and returned to a graded series of cryoprotectant solution diluted in PBS. Sections were then submerged in a preincubation solution prepared with 1% normal goat serum (NGS) and 1% bovine serum albumin (BSA) in PBS at RT for 60 min. Following this, sections were placed in the primary antibody solution at RT for 24 hr. The primary antibody solution was prepared with specific antibodies directed against either vGluT1 (guinea pig, RRID: AB_2301751, dilution 1:5,000) or vGluT2 (rabbit, RRID: AB_2315568, dilution 1:5,000), diluted in PBS containing 1% NGS and 1% BSA. After the primary antibody incubation, sections were thoroughly rinsed in PBS and placed for 90 min at RT in a secondary antibody solution consisting of 1% NGS, 1% BSA and the secondary biotinylated antibody raised against the primary antibodies, a goat anti-guinea pig IgG (Cat# BA-7000, Vector, dilution 1:200) for vGluT1 and a goat anti-rabbit IgG (Cat# BA-1000, Vector, dilution 1:200) for vGluT2. Afterwards, sections were thoroughly rinsed in PBS and placed at RT for 90 min in an avidin-biotinylated peroxidase complex (ABC; Cat# PK-4000, Vector Laboratories) diluted 1:100 in PBS containing 1% BSA. After incubation, sections were rinsed in PBS and TRIS buffer (0.05M pH 7.6) before being placed in a solution containing 0.025% diaminobenzidine (3,30-diaminobenzidine tetrahydrochloride, DAB; Sigma, St Louis, MO), 0.01 M imidazole (Fisher Scientific, Norcross, GA) and 0.005% H_2_O_2_ for 10 min at RT. The reaction was stopped by several washes in PBS.

#### 2.3.2 Tissue processing for transmission electron microscopy (TEM) and image acquisition

Following the DAB reaction, sections were transferred to a phosphate buffer solution (PB, 0.1M, pH 7.4). The tissue was then postfixed in a 1% osmium tetroxide solution for 20 min. After washes in PB, sections were dehydrated through a graded series of alcohol solutions (50-100%) before placed in propylene oxide. To enhance contrast in the EM, uranyl acetate (1%) was added to the 70% alcohol solution during dehydration. Following dehydration, sections were embedded in epoxy resin (Durcupan ACM; Fluka, Buchs, Switzerland) for 12 hr, mounted onto oil-coated slides, coverslipped, and polymerized in the oven at 60℃ for 48 hr. Square blocks of dorsal striatum tissue were cut out from the slides and glued onto resin blocks. Serial ultrathin sections (60 nm-thickness) were obtained using an ultramicrotome (Ultracut T2; Leica, Germany), collected onto Pioloform coated single-slot copper grids, stained with lead citrate (5 min), and examined with a TEM (JEOL/JEM-1011). Images were acquired using a CCD Camera (Gatan Model 785).

#### 2.3.3 Tissue processing and image acquisition for AT/SEM

After stopping the DAB reaction with PBS washes, circular punches of tissue (1-mm diameter) were taken from the dorsolateral striatum of immunostained sections and shipped to the Oregon Health Science Center University Microscopy Core (Portland, Oregon) in 4% paraformaldehyde for array tomography/scanning electron microscopy (AT/SEM) processing. Samples were first rinsed in 0.1M sodium cacodylate buffer and stained with tannic acid (1%) in 0.1 M sodium cacodylate for 15 min. Tissue was then incubated in osmium tetroxide (2%) reduced with potassium ferricyanide (1.5%) in 0.1 M sodium cacodylate for 1 hr, followed by overnight incubation in uranyl acetate (1%) at 4 °C. Then, samples were treated with lead aspartate at 60 °C for 1 hr, dehydrated through acetone series (50%, 75% 85%, 95%, 100%), and infiltrated with resin overnight. Finally, samples were further infiltrated with resin and polymerized in the oven at 60℃. Ultrathin sections (70 nm) were cut using a Leica Artos 3D ultramicrotome equipped with a Leica AT-4 diamond knife and collected as ribbons onto silicon chips. The chips were mounted on a stub and imaged using a Helios 5 UC scanning electron microscope (Thermo Fisher Scientific) with Maps™ software array tomography module. Regions of interest were acquired at 4-nm pixel resolution (10,240 x 10,240 pixels) using the CBS detector. Image stacks (∼150 to 200 serial images per stack) were aligned in FIJI using the *Linear Stack Alignment with SIFT* plugin, with the expected transformation set to affine. The aligned images were then imported into Amira (Thermo Fisher Scientific), to extract a 6144 x 6144 pixels subvolume.

### 2.4 Data analysis

To ensure that all electron microscopic analyses were performed in an unbiased manner, the experimenter responsible for data collection and analysis was blinded to the experimental group from which the tissue originated.

#### 2.4.1 Postsynaptic targets of vGluT1- and vGluT2-IR terminals: TEM

For TEM, a total of six blocks of vGluT1-immunostained and six blocks of vGluT2-immunostained striatal tissue were analyzed, with one block of each type obtained from each animal (N = 6) and data collected from 1–4 grids per block. Approximately 50–80 electron micrographs of randomly distributed vGluT1- and vGluT2-IR terminals were acquired per animal at 40,000X magnification from superficial regions of the tissue to ensure optimal antibody penetration (Table 3). Terminals were identified by the presence of a dark, amorphous DAB peroxidase deposit, and the formation of clear synaptic contacts with postsynaptic dendritic spines or shafts. Images were analyzed using Gatan Digital Micrograph software, and vGluT1- and vGluT2-IR terminals were identified and counted.

**Table 3:** Number of axo-spinous and axo-dendritic synapses.

| **vGluT1** |  |  |  |
| --- | --- | --- | --- |
| **Number of Synapses** | **Axo-spinous** | **Axo-dendritic** | **Total images** |
| WT (RM-32) | 54 | 2 | 56 |
| WT (RM-166) | 83 | 0 | 83 |
| WT (RM-172) | 79 | 7 | 86 |
| KO (RM-31) | 64 | 2 | 66 |
| KO (RM-175) | 70 | 6 | 76 |
| KO (RM-177) | 73 | 3 | 76 |

| **vGluT2** |  |  |  |
| --- | --- | --- | --- |
| **Number of Synapses** | **Axo-spinous** | **Axo-dendritic** | **Total images** |
| WT (RM-32) | 42 | 5 | 47 |
| WT (RM-166) | 54 | 7 | 61 |
| WT (RM-172) | 58 | 7 | 65 |
| KO (RM-31) | 39 | 12 | 51 |
| KO (RM-175) | 45 | 20 | 65 |
| KO (RM-177) | 67 | 14 | 81 |

For each terminal, the postsynaptic target (dendritic spine or shaft) was determined, and synapses were classified as axo-spinous or axo-dendritic accordingly. Spines were defined as protrusions emerging from dendritic shafts. Because data were collected from single ultrathin sections, the parent dendrite from which spines originate from was not always visible; additional ultrastructural criteria, including the presence of a spine apparatus and absence of mitochondria, were used to differentiate dendritic spine profiles from small dendritic shafts. The percentage of axo-spinous and axo-dendritic synapses formed by vGluT1- and vGluT2-IR terminals was calculated for each animal by dividing the number of synapses in each category by the total number of corresponding vGluT1- and vGluT2-IR terminals analyzed. Group data are presented as mean percentage ± SEM, where each data point represents an individual animal (N = 3) and error bars indicate variability between animals.

#### 2.4.2 3D ultrastructural analysis of vGluT1- and vGluT2-IR synapses

Using an AT/SEM, serial digitized electron microscopic images (100–200 per series/stack; TIFF format) of vGluT1- or vGluT2-IR striatal terminals from three WT and three KO mice were imported into the *Reconstruct* [NIH] software (https://synapseweb.clm.utexas.edu/software-0) for 3D reconstruction and quantitative ultrastructural comparative analysis. After images were imported in *Reconstruct* and before element segmentation and tracing, section thickness (70 nm) and pixel size (0.004 µm) were specified in the software. Immunoreactive terminals were identified and selected based on the presence of a dark, amorphous DAB peroxidase deposit and the formation of asymmetric synapses with dendritic spines or shafts. In each serial section, presynaptic terminals, postsynaptic targets (dendritic spines and shafts), and PSDs were manually traced. The software then generated complete 3D reconstructions of individual synapses. The following ultrastructural parameters were quantified and compared between WT and KO mice: (1) presynaptic terminal volume (µm^3^), (2) dendritic spine volume (µm^3^), and (3) PSD area (µm^2^). Additionally, PSDs were classified as either macular or perforated in vGluT1-IR axo-spinous synapses and vGluT2-IR axo-spinous and axo-dendritic synapses. Macular PSDs were defined as continuous, non-segmented densities, whereas perforated PSDs were characterized by discontinuous, segmented densities, as described in previous studies (Geinisman, 1993; Harris, 2020; Harris & Weinberg, 2012; Neuhoff et al., 1999).

### 2.5 Statistical analysis

Interindividual variability among animals within each group for TEM analyses was evaluated using the Kruskal-Wallis test (GraphPad Prism 11). Statistical comparisons between WT and KO mice for TEM data were performed using Welch’s t-test (GraphPad Prism 11). Interindividual variability among animals within each group for 3D reconstruction analyses was assessed for each ultrastructural parameter using one-way ANOVA, along with Bartlett’s and Brown–Forsythe tests to evaluate homogeneity of variance (GraphPad Prism 11). Some parameters exhibited variability between animals and/or heterogeneity of variance, indicating that variability was parameter dependent. Accordingly, all comparisons between WT and KO animals were performed using nonparametric tests, with the animal as the unit of analysis. Ultrastructural comparisons were assessed using the Wilcoxon rank-sum test (GraphPad Prism 11) due to non-normal data distribution. PSD morphology comparisons between WT and KO mice were evaluated using Welch’s t-test (GraphPad Prism 11). All data are presented as mean ± SEM, with error bars representing variability between animals.

### 2.6 Photographs Production

The electron micrographs shown in this manuscript were digitally acquired, imported in TIFF format to Adobe Photoshop (CC 2019; Adobe Systems, San Jose, CA), and adjusted only for brightness and contrast to optimize the quality of the images for analysis. Micrographs were then compiled into figures using Adobe Photoshop CC.

## 3 RESULTS

### 3.1 Quantitative analysis of corticostriatal and thalamostriatal synapses using single TEM images

Previous studies reported a reduction in thalamostriatal, but not corticostriatal, excitatory input in mice with striatal-specific ablation of GluD1 (Liu et al., 2020). Because the postsynaptic targets of synapses determine how excitatory inputs are integrated within dendrites, analyses of synaptic connectivity patterns provide important insight into circuit organization (Raju et al., 2006). To assess potential alterations in excitatory synaptic organization, we analyzed the ultrastructural features and synaptic connectivity patterns of corticostriatal (vGluT1-IR) and thalamostriatal (vGluT2-IR) terminals in the dorsolateral striatum of WT and KO mice. For each terminal, the postsynaptic target (dendritic spine or shaft) was quantified (Figures 1 and 2; Table 3).

**Figure 1:**
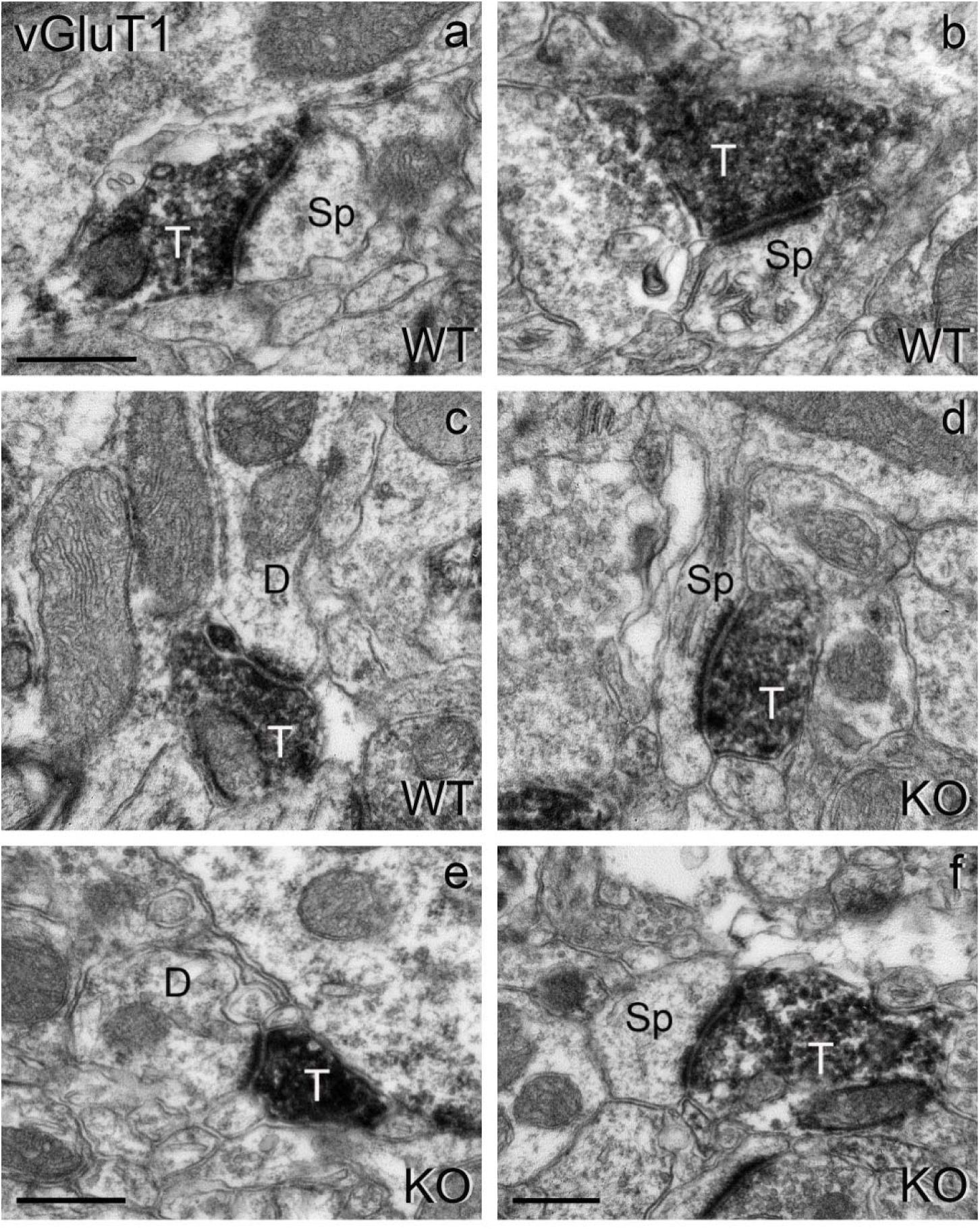
Electron micrographs (TEM) of vGluT1-IR terminals in the mouse striatum. vGluT1-IR terminals form asymmetric synapses with dendritic spines (a, b, d, and f) and dendritic shafts (c and e) in WT (a–c) and KO (d–f) mice. T: terminal; Sp: dendritic spine; D: dendrite. Scale bar in (a) applies to (b–d). Scale bars = 0.4 µm in all panels.

**Figure 2:**
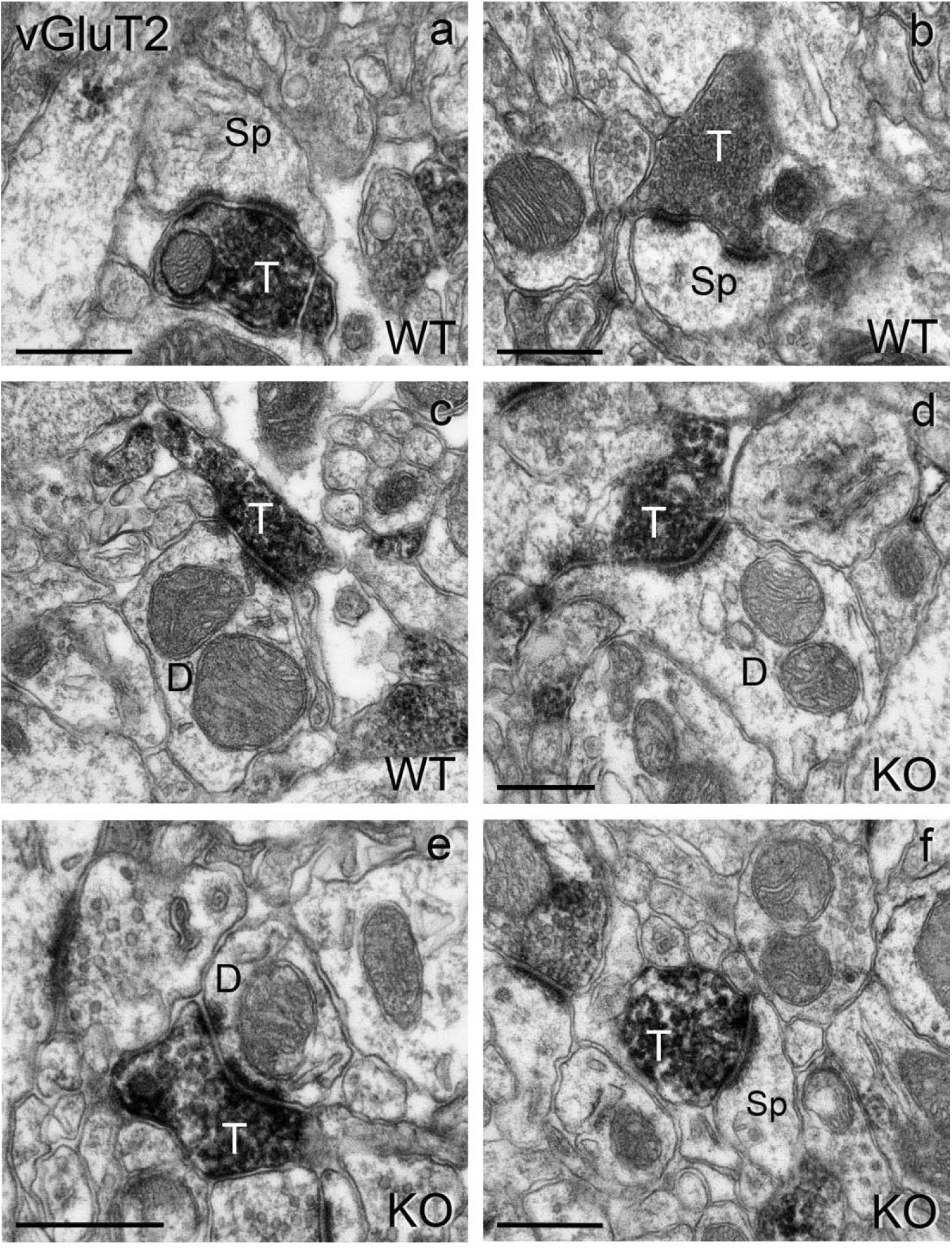
Electron micrographs (TEM) of vGluT2-IR terminals in the mouse striatum. vGluT2-IR terminals form asymmetric synapses with dendritic spines (a, b, and f) and dendritic shafts (c–e) in WT (a–c) and KO (d–f) mice. T: terminal; Sp: dendritic spine; D: dendrite. Scale bar in (b) applies to (c). Scale bars = 0.4 µm in all panels.

### Corticostriatal axon terminals

A total of 443 vGluT1-IR corticostriatal terminals and their postsynaptic elements were analyzed in the dorsolateral striatum of WT (n = 225) and KO (n = 218) mice (Figure 1). These vGluT1-IR terminals formed asymmetric synapses with either dendritic spines (Figure 1a,b,f) or dendritic shafts (Figure 1c–e) in both WT (Figure 1a–c) and KO (Figure 1d–f) mice. In WT mice, 96.1 ± 2.4% (mean ± SEM; N = 3) of vGluT1-IR terminals formed synapses with dendritic spines, whereas 3.9 ± 2.4% targeted dendritic shafts (Figure 3a). Axo-spinous synapses were significantly more prevalent than axo-dendritic synapses (paired t-test, **p = 0.0026). No significant difference in the relative percentages of axo-spinous and axo-dendritic synapses was found between WT and KO mice (Welch’s t-test, p = 0.728) (Figure 3a).

**Figure 3:**
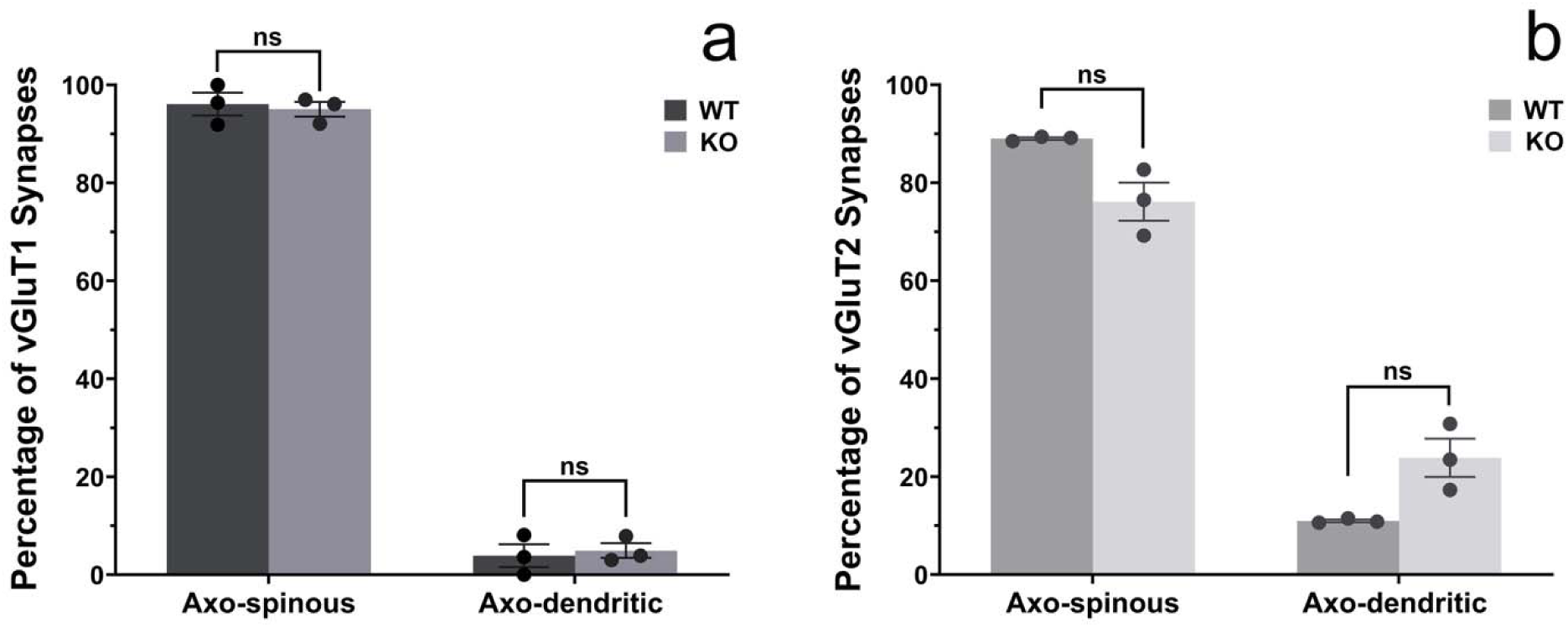
Bar graphs comparing the percentages of axo-spinous and axo-dendritic synapses formed by vGluT1-IR (a) and vGluT2-IR (b) terminals in the striatum of WT and KO mice. (a) In WT mice, 96.1 ± 2.4% of the vGluT1-IR terminals (n = 225) form axo-spinous synapses, while 3.9 ± 2.4% form axo-dendritic. In KO mice, 95.0 ± 1.5% of the vGluT1-IR terminals (n = 218) form axo-spinous synapses while 5.0 ± 1.5% form axo-dendritic. No significant difference (Welch’s t-test, p = 0.73) was found in the distribution of axo-spinous and axo-dendritic synapses of vGluT1-IR terminals between WT and KO mice (N = 3/group). (b) In WT mice, 89.0 ± 0.26% of the vGluT2-IR terminals (n = 173) form axo-spinous synapses while 11.0 ± 0.26% form axo-dendritic. In KO mice, 76.1 ± 3.9% of the vGluT2-IR terminals (n = 197) form axo-spinous synapses, while 23.9 ± 3.9% form axo-dendritic. No significant difference (Welch’s t-test, p = 0.08) was found in the distribution of synaptic targets of thalamostriatal vGluT2-IR terminals between WT and KO mice.

### Thalamostriatal axon terminals

A total of 370 vGluT2-IR thalamostriatal terminals and their postsynaptic elements were analyzed in the dorsolateral striatum of WT (n = 173) and KO (n = 197) mice (Figure 2). These vGluT2-IR terminals formed asymmetric synapses with either dendritic spines (Figure 2a,b,f) or dendritic shafts (Figure 2c–e) in both WT (Figure 2a–c) and KO (Figure 2d–f) mice. In WT mice, 89.0 ± 0.26% of vGluT2-IR terminals formed synapses with dendritic spines, while 11.0 ± 0.26% targeted dendritic shafts (Figure 3b). As observed for corticostriatal terminals, axo-spinous synapses were significantly more prevalent than axo-dendritic synapses (paired t-test, ****p < 0.0001). When compared with KO mice, no significant difference in the relative percentages was found (Welch’s t-test, p = 0.0798) (Figure 3b).

### 3.2 Ultrastructural 3D analysis of corticostriatal and thalamostriatal synapses

To further investigate the synaptic ultrastructure of corticostriatal and thalamostriatal synapses, serial EM images acquired via AT/SEM and *Reconstruct* [NIH] software were used to generate complete 3D models and quantitative morphometric data. Terminals were identified by the presence of the dark, amorphous DAB deposit (Figure 4a1–3,c1–3, 5a1–3,c1–3, and 6a1–3,c1–3) that formed asymmetric synapses with either dendritic spines or shafts (Table 4). Synaptic contacts were identified based on the aggregation of synaptic vesicles at the presynaptic terminal and the thickening of the postsynaptic membrane.

**Figure 4:**
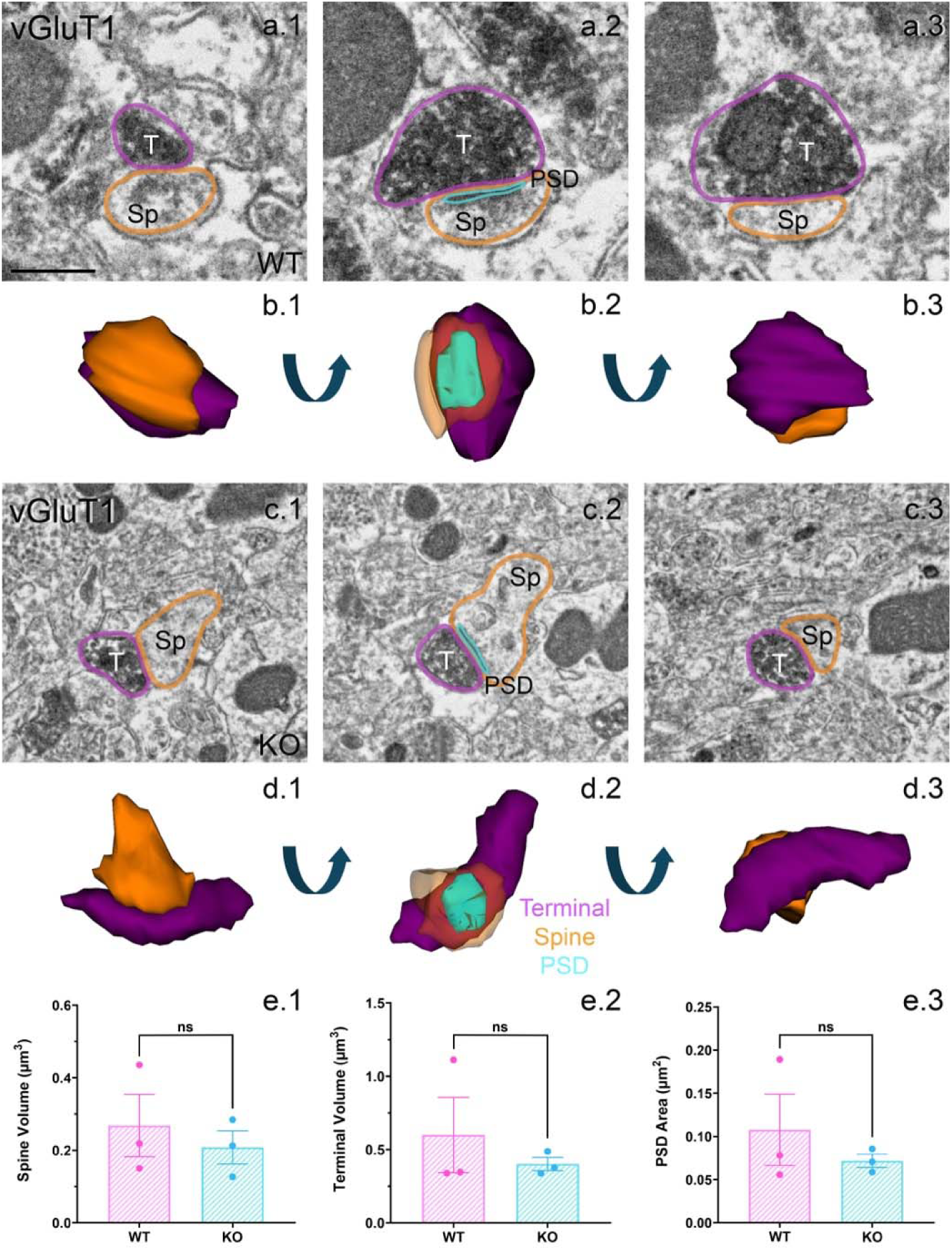
3D reconstruction of axo-spinous synapses formed by vGluT1-IR terminals in the mouse striatum. (a1–3, c1–3) Serial EM images of an axo-spinous synapse in the striatum of a WT (a1–3) and a KO (c1–3) mouse. (a1–3) Images of EM sections 64, 68, and 72, respectively; 19 consecutive EM images were used for the 3D reconstruction of this macular asymmetric synapse (b1–3). (c1–3) Images of EM sections 20, 24, and 27, respectively; 24 consecutive EM images were used for the 3D reconstruction of this macular asymmetric axo-spinous synapse (d1–3). For the 3D reconstructions, the elements of the synapses were segmented and traced in every consecutive image using *Reconstruct.* The models were rotated to allow different views of the synaptic components. (e1–3) Histograms comparing the spine volume (e1), terminal volume (e2), and PSD area (e3) with bar graphs comparing WT (N = 3) and KO (N = 3) mice. No significant differences were found between the two groups (Wilcoxon rank-sum test: spine volume, p = 0.70; terminal volume, p > 0.99; PSD area, p > 0.99). T: terminal; Sp: dendritic spine; PSD: postsynaptic density. Scale bar in (a1) applies to (a2, a3, and c1–3) = 0.25 µm.

**Table 4:** Number of synapses reconstructed.

| Animal | vGluT1 | vGluT2 |  |
| --- | --- | --- | --- |
|  | axo-spinous synapses | axo-spinous synapses | axo-dendritic synapses |
| WT (RM-172) | 10 | 10 | 11 |
| WT (RM-202) | 10 | 10 | 9 |
| WT (RM-204) | 10 | 10 | 8 |
| KO (RM-171) | 11 | 13 | 7 |
| KO (RM-177) | 10 | 10 | 11 |
| KO (RM-206) | 10 | 11 | 11 |

Specifically, vGluT1-IR (corticostriatal) and vGluT2-IR (thalamostriatal) glutamatergic synapses were analyzed in the dorsolateral striatum of WT (N = 3) and KO (N = 3) mice. In WT mice, 30 vGluT1 axo-spinous, 30 vGluT2 axo-spinous, and 28 vGluT2 axo-dendritic terminals, along with their corresponding postsynaptic target(s) and PSDs, were fully reconstructed and morphometrically analyzed. In KO mice, 31 vGluT1 axo-spinous, 34 vGluT2 axo-spinous, and 29 vGluT2 axo-dendritic terminals and their synapses were similarly reconstructed and analyzed. Because dendritic shafts could not be reliably reconstructed in their entirety from serial EM stacks, dendritic shaft morphometry was not analyzed; size measurements would likely reflect sectioning orientation and truncation rather than true morphological variation between WT and KO mice. Quantitative data reported in Figures 4–6 represent the mean values ± SEM from three mice per genotype.

**Figure 5:**
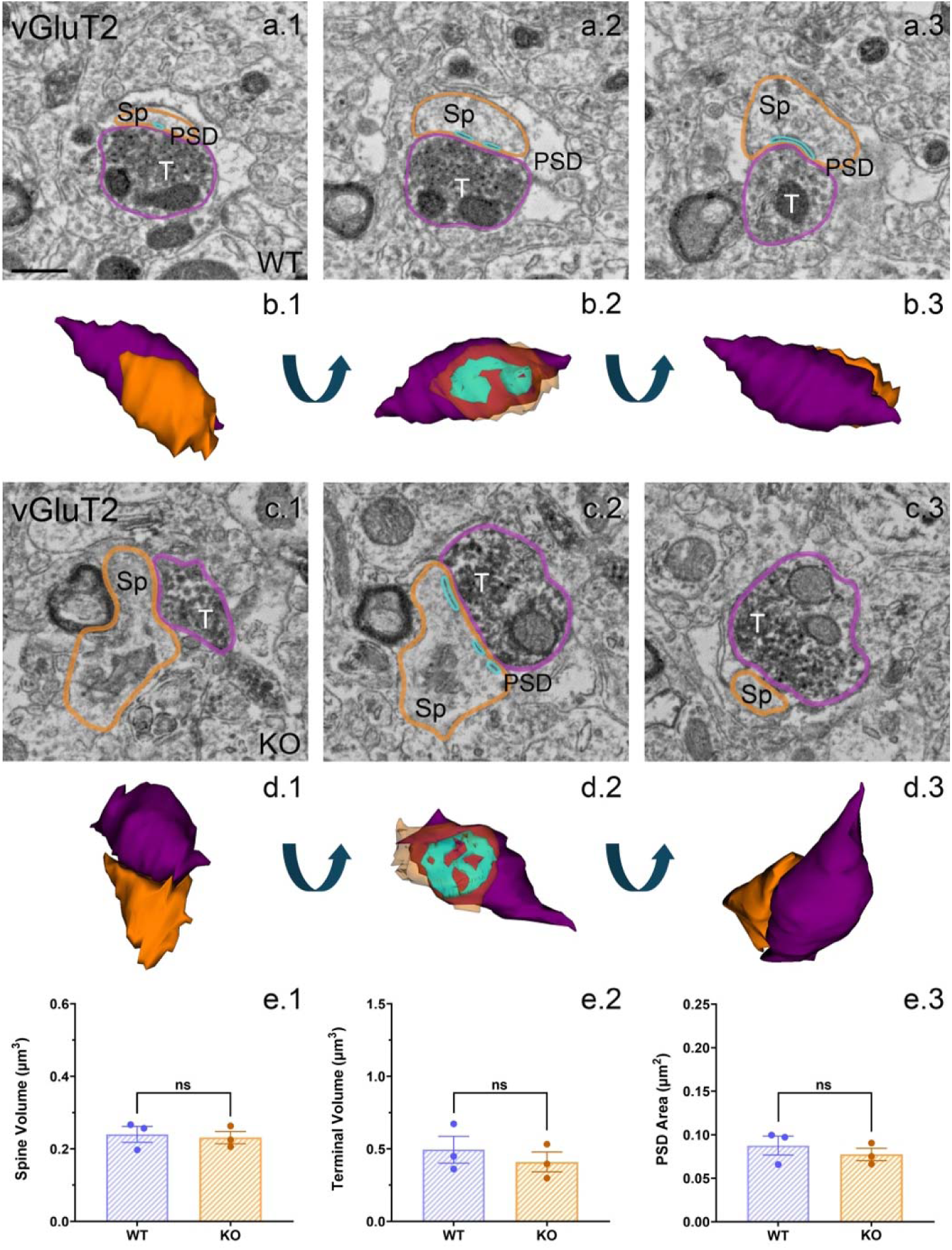
3D reconstruction of axo-spinous synapses formed by vGluT2-IR terminals in the mouse striatum. (a1–3, c1–3) Serial EM images of an axo-spinous synapse in the striatum of a WT (a1–3) and KO (c1–3) mouse. (a1–3) Images of EM sections 18, 24, and 29, respectively; 34 consecutive EM images were used for the total reconstruction of this perforated asymmetric axo-spinous synapse (c1–3). Images of EM sections 39, 43, and 52, respectively; 31 consecutive EM images were used for the total reconstruction of this asymmetric perforated axo-spinous synapse. For the 3D reconstructions, the elements of the synapses were segmented and traced in every consecutive image using *Reconstruct.* The models were rotated to allow different views of the synaptic components. (e1–3) Histograms comparing the spine volume (e1), terminal volume (e2), and PSD area (e3) with bar graphs comparing WT (N = 3) and KO (N = 3) mice. No significant differences were found between the two groups (Wilcoxon rank-sum test: spine volume, p > 0.99; terminal volume, p = 0.70; PSD area, p = 0.70). T: terminal; Sp: dendritic spine; PSD: postsynaptic density. Scale bar in (a1) applies to (a2, a3, and c1–3) = 0.25 µm.

**Figure 6:**
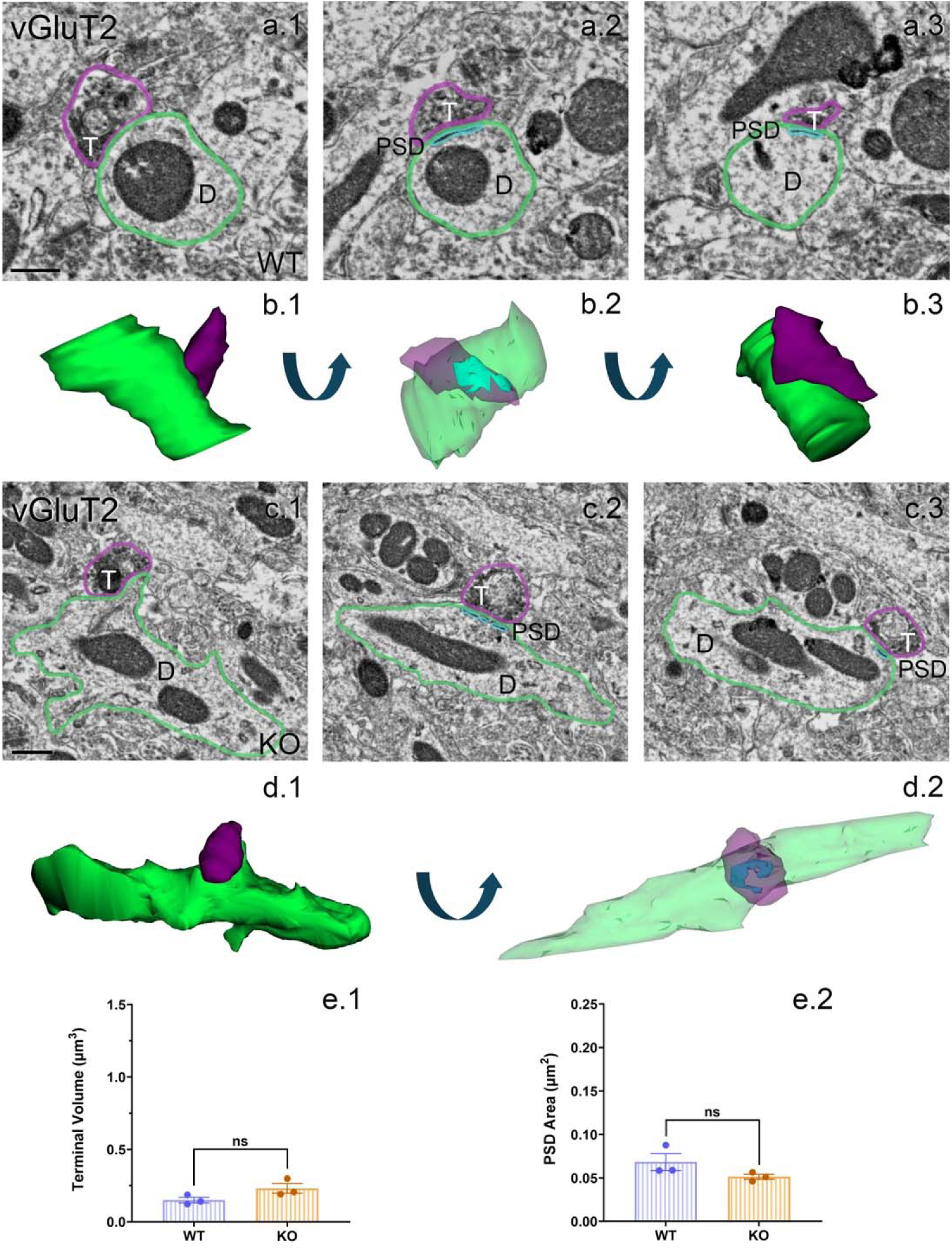
3D reconstruction of axo-dendritic synapses formed by vGluT2-IR terminals in the mouse striatum. (a1–3, c1–3) Serial EM images of an axo-dendritic synapse in the striatum of a WT (a1–3) and a KO (c1–3) mouse. (a1–3) Images of EM sections 44, 48, and 53, respectively; 22 consecutive EM images were used for the total reconstruction of this perforated asymmetric axo-dendritic synapse (b1-3). (c1–3) Images of EM sections 6, 9, and 13, respectively; 20 consecutive EM images were used for the total reconstruction of this perforated asymmetric axo-dendritic synapse (d1–3). For the 3D reconstructions, the elements of the synapses were segmented and traced in every consecutive image using *Reconstruct.* The models were rotated to allow different views of the synaptic components. (e1–2) Histograms comparing the terminal volume (e1) and PSD area (e2) with bar graphs comparing WT (N = 3) and KO (N = 3) mice. No significant differences were found between the two groups (Wilcoxon rank-sum test: terminal volume, p = 0.10; PSD area, p = 0.10). T: terminal; D: dendrite; PSD: postsynaptic density. Scale bar in (a1) applies to (a2, a3). Scale bar in (c1) applies to (c2, c3). Scale bars = 0.25 µm in all panels.

PSD morphology was also assessed using 3D reconstructed models (Figure 7). Reconstruction across serial sections allows for the visualization of the full PSD architecture, ensuring reliable identification of PSD morphology as either perforated or macular. In single 2D sections, perforated PSDs may appear to have a continuous density if the discontinuous segments are not captured within a given section.

**Figure 7:**
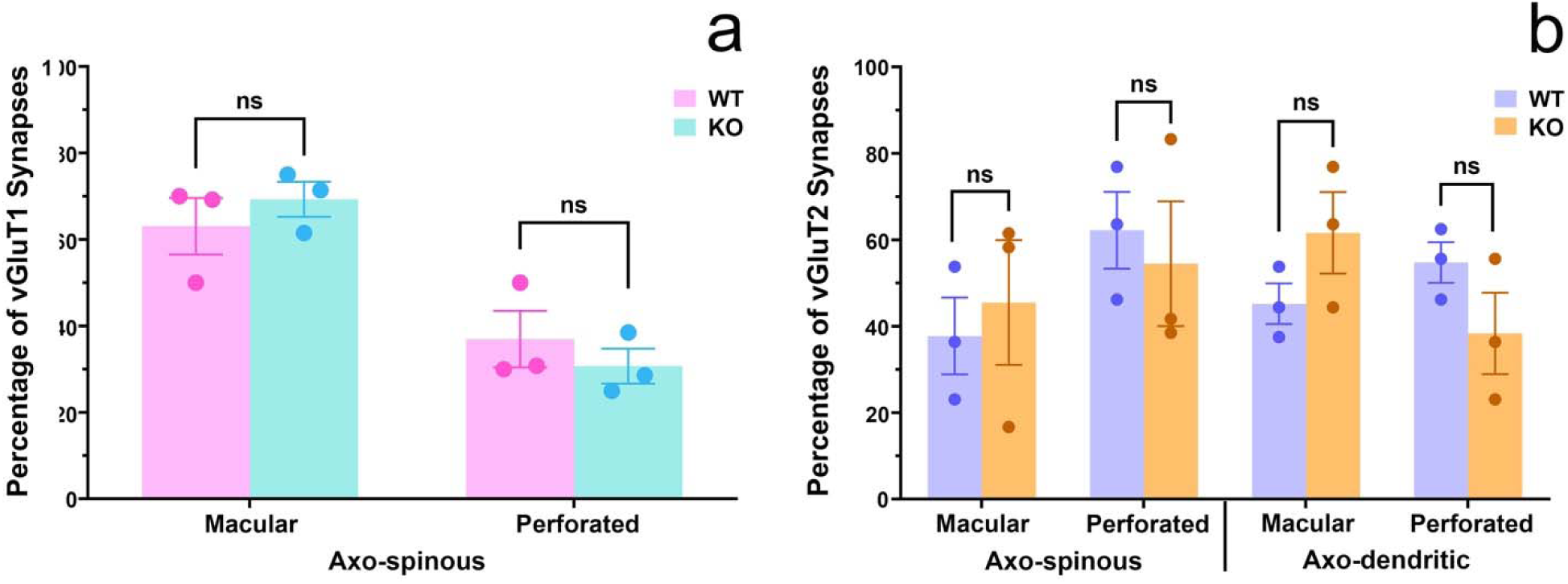
Histograms showing the comparative percentages of macular and perforated PSDs in synapses formed by vGluT1- and vGluT2-IR terminals in the striatum of WT and KO mice. (a) For vGluT1-IR terminals, the percentage of macular versus perforated PSDs in axo-spinous synapses was not significantly different between WT (macular: 63.1 ± 6.5%; perforated: 36.9 ± 6.5%) and KO (macular: 69.3 ± 4.0%; perforated: 30.7 ± 4.0%) mice (Welch’s t-test, p = 0.47). (b) Similarly, for vGluT2-IR terminals, the percentage of macular versus perforated PSDs in axo-spinous synapses was not significantly different between WT (macular: 37.8 ± 6.5%; perforated: 62.2 ± 6.5%) and KO (macular: 45.5 ± 14%; perforated: 54.5 ± 14%) mice. Likewise, the percentage of macular versus perforated PSDs in axo-dendritic synapses was not significantly different between WT (macular: 45.2 ± 4.7%; perforated: 54.8 ± 4.7%) and GluD1-KO (macular: 61.6 ± 9.4%; perforated: 38.4 ± 9.4%) mice (Welch’s t-test: axo-spinous, p = 0.68; axo-dendritic, p = 0.22).

#### 3.2.1 Corticostriatal synapses

##### Spine volume

A total of 68 dendritic spines contacted by vGluT1-IR corticostriatal terminals were reconstructed using serial images in WT (n = 33) and KO (n = 35) mice. The mean spine volume was 0.27 ± 0.086 µm³ in WT mice and 0.21 ± 0.045 µm³ in KO mice. Comparative quantitative analysis showed no significant difference between WT and KO mice (two-sided Wilcoxon rank sum test, p = 0.70) (Figure 4e1 and 1Sa).

##### Terminal volume

Morphometric analysis was performed on a total of 61 vGluT1-IR terminals forming axo-spinous synapses from WT (n = 30) and KO (n = 31) mice (Table 4). The number of vGluT1-IR terminals is less than the number of spines because some individual terminals contacted multiple spines. In WT mice, the mean terminal volume was 0.60 ± 0.26 µm^3^. Although it was slightly smaller (0.40 ± 0.045 µm³) in KO mice, comparative quantitative analysis showed no significant difference between WT and KO mice (two-sided Wilcoxon rank-sum test, p > 0.99) (Figure 4e2 and 1Sb).

##### PSD area and morphology

For these synapses, the mean PSD area was also not significantly different between WT (0.11 ± 0.041 µm²) and KO (0.072 ± 0.0078 µm²) mice (two-sided Wilcoxon rank-sum test, p > 0.99) (Figure 4e3 and 1Sc). Additionally, the proportion of macular versus perforated PSDs did not differ significantly between the two groups (WT = 63.1 ± 6.5% macular; KO = 69.3 ± 4.0% macular; Welch’s t-test, p = 0.47) (Figure 7a and 1Sc).

#### 3.2.2 Thalamostriatal synapses

##### Spine volume

A total of 74 dendritic spines contacted by vGluT2-IR thalamostriatal terminals were reconstructed using serial images in WT (n = 37) and KO (n = 37) mice. The mean spine volume was 0.24 ± 0.022 µm^3^ in WT and 0.23 ± 0.017 µm³ in KO mice. No significant difference was observed between WT and KO mice (two-sided Wilcoxon rank-sum test, p > 0.99) (Figure 5e1 and 2Sa).

##### Terminal volume

A total of 121 vGluT2-IR terminals were reconstructed in WT and KO mice (Table 4). Of these terminals, 64 formed synapses with dendritic spines in WT (n = 30) and KO (n = 34) mice (Figure 5), whereas 57 formed synapses directly on the dendritic shaft in WT (n = 28) and KO (n = 29) mice (Figure 6).

For axo-spinous thalamostriatal synapses, no significant difference was found in the mean terminal volume between WT (0.49 ± 0.093 µm^3^) and KO (0.41 ± 0.068 µm^3^) mice (two-sided Wilcoxon rank-sum test, p = 0.70) (Figure 5e2 and 2Sb).

For axo-dendritic thalamostriatal synapses, the mean terminal volume in KO (0.23 ± 0.034 µm^3^) was larger than in WT (0.15 ± 0.019 µm^3^) mice, but the difference was not statistically significant (two-sided Wilcoxon rank-sum test, p = 0.10) (Figure 6e1 and 2Sd).

Differences in the terminal volume of axo-spinous and axo-dendritic synapses were also compared in WT and KO mice. In both groups, no significant differences were found; however, terminals forming axo-spinous synapses were slightly larger than those forming axo-dendritic synapses (WT: p = 0.25; KO: p = 0.50; Wilcoxon signed-rank test) (Figure 3Sa,c).

##### PSD area and morphology

For the axo-spinous synapses, the mean PSD area in WT (0.088 ± 0.011 µm^2^) was closely similar to that in KO (0.077 ± 0.0071 µm^2^) mice, and the difference was not significant (two-sided Wilcoxon rank-sum test, p = 0.70) (Figure 5e3 and 2Sc).

Similarly, the mean PSD area of axo-dendritic synapses in WT (0.068 ± 0.0097 µm^2^) and KO (0.051 ± 0.0029 µm^2^) mice were not significantly different (two-sided Wilcoxon rank-sum test, p = 0.10) (Figure 6e2 and 2Se).

Furthermore, the proportions of macular versus perforated PSDs associated with vGluT2-IR axo-spinous and axo-dendritic synapses showed no significant differences between WT and KO mice (axo-spinous: WT = 37.8 ± 6.5% macular, KO = 45.5 ± 14% macular, p = 0.68; axo-dendritic: WT = 45.2 ± 4.7% macular, KO = 61.6 ± 9.4% macular, p = 0.22; Welch’s t-test) (Figure 7b). Differences in the PSD area of axo-spinous and axo-dendritic synapses were also compared in WT and KO mice. In both groups, no significant differences were found (WT: p = 0.25; KO: p = 0.25; Wilcoxon signed-rank test) (Figure 3Sb,d).

## 4 DISCUSSION

The present study examined synaptic connectivity patterns and ultrastructural features of corticostriatal and thalamostriatal glutamatergic terminals and their synapses in the dorsolateral striatum of GluD1 KO mice. Using TEM, we found that both vGluT1- and vGluT2-IR terminals formed a greater proportion of synapses onto dendritic spines than onto dendritic shafts, and that this overall pattern of synaptic targeting was preserved in KO mice. Given our previous observation that GluD1 immunoreactivity is preferentially expressed in striatal dendritic shafts relative to spines (Hoover et al., 2020; Liu et al., 2020), these findings suggest that GluD1 localization and subcellular innervation patterns may not be directly coupled. The enrichment of striatal GluD1 within dendritic shafts may reflect roles beyond regulating thalamic afferents, consistent with previous findings of GluD1 immunoreactivity at nonsynaptic neuronal plasma membranes, in glia, and on mitochondria outer membranes (Hoover et al., 2020; Liu et al., 2020). To further investigate the role of GluD1 in synaptic organization, we used 3D reconstructions to achieve a more complete characterization of synaptic architecture. In contrast to 2D analyses, which are constrained by the sectioning plane, this approach captures the full morphology of postsynaptic spines, presynaptic terminals, and PSDs. Quantitative analysis of reconstructed corticostriatal and thalamostriatal axo-spinous synapses revealed no significant morphometric differences between KO and WT mice in spine volume, terminal volume, or PSD area. Similarly, analysis of reconstructed thalamostriatal axo-dendritic synapses showed no major differences between the two genotypes in terminal volume or PSD area.

### 4.1 Preservation of synaptic ultrastructure despite reduced thalamostriatal PF-MSN input

Previous studies have shown that striatum-specific ablation of GluD1 in mice leads to a significant reduction in miniature excitatory postsynaptic current (mEPSC) frequency in striatal medium spiny neurons (MSNs), accompanied by a selective decrease in thalamostriatal vGluT2-IR input, while corticostriatal vGluT1-IR input remained unchanged (Liu et al., 2020). Because circuit-level function is closely linked to both patterns of synaptic connectivity (Raju et al., 2006) and the ultrastructure of synaptic components (Harris et al., 1992; Harris & Kater, 1994; Holler et al., 2021; Pan et al., 2023; Villalba & Smith, 2010), we examined whether these functional deficits could be, in part, explained by morphometric changes at the level of individual synapses. In addition, through 3D reconstruction of complete PSDs, we assessed potential alterations in their morphology, commonly classified as macular or perforated. PSD morphology is widely regarded as an ultrastructural correlate of synaptic strength and plasticity, with perforated PSDs often associated with enhanced synaptic efficacy (Borczyk et al., 2019; Bourne & Harris, 2008; Geinisman, 1993, 2000; Harris, 2020; Harris et al., 2003; Lisman, 2017; Lisman & Harris, 1993; Luscher et al., 2000; Neuhoff et al., 1999; Sorra & Harris, 2000). Specifically, the perforations and compartmentalized structure of perforated PSDs support multiple transmission zones and reduce postsynaptic receptor saturation (Bell et al., 2014; Chirillo et al., 2019; Harris, 2020; Lisman & Harris, 1993). Our findings indicate that, despite reduced thalamostriatal input (Liu et al., 2020), synaptic targeting and ultrastructural features of vGluT2-IR thalamostriatal synapses are preserved in KO mice. The absence of structural abnormalities at individual synaptic contacts suggests that the reduction in excitatory transmission is driven by a decreased density of thalamic terminals, rather than by morphological alterations that would attenuate the potency of single synapses. Furthermore, the absence of detectable structural alterations in the remaining synapses suggests a lack of compensatory ultrastructural remodeling in response to reduced input.

### 4.2 GluD2 as a model for GluD1 function in synaptogenesis

While the role of GluD1 in the central nervous system (CNS) remains incompletely understood, its family member GluD2 has been extensively studied, particularly in the cerebellum. Previous studies have shown that GluD2 is critical for the formation of parallel fiber-Purkinje cell synapses, as GluD2-KO mice exhibit normal neuronal and spine morphology but a reduced number of synapses, indicating a specific role in synaptogenesis (Kurihara et al., 1997). In addition, GluD2 functions as a synaptic assembly molecule, enabling synapse regeneration following parallel fiber transection (Ichikawa et al., 2016). In this context, prior findings demonstrating a reduction in thalamostriatal projections in mice with striatum-specific GluD1 ablation (Liu et al., 2020), together with our observation that the ultrastructural features of remaining vGluT2-IR thalamostriatal synapses are preserved, suggest that GluD1 plays a similar role in the striatum by contributing to the establishment or maintenance of specific synaptic inputs, particularly thalamostriatal projections, without being essential for regulating synaptic ultrastructure. At the molecular level, GluD1 forms trans-synaptic adhesion complexes with presynaptic neurexins via cerebellin 1 (Cbln1), which is highly expressed in Pf thalamic neurons (Kusnoor et al., 2010; Miura et al., 2006; Otsuka et al., 2016). Because Pf neurons are a principal source of glutamatergic axo-dendritic synapses onto MSNs (Dube et al., 1988; Sadikot et al., 1992), these interactions are well positioned to regulate the formation and stabilization of Pf-MSN synapses. Loss of GluD1 would therefore be expected to disrupt these trans-synaptic interactions. Given the preservation of ultrastructural features in vGluT2-IR thalamostriatal axo-dendritic synapses in KO mice, our 3D analysis suggests that not all Pf-MSN synapses are disrupted in the absence of GluD1. It is likely that thalamostriatal inputs comprise distinct subpopulations: a GluD1-dependent population that is lost and a GluD1-independent population that remains structurally intact.

### 4.3 Other potential mechanisms in GluD1-mediated synaptic regulation

The preservation of synaptic ultrastructure in surviving vGluT2-IR thalamostriatal synapses supports the idea that the reduced thalamostriatal projections observed is the primary contributor to the observed physiological reduction in thalamic input in mice with striatal-specific GluD1 ablation (Liu et al., 2020). However, because our 3D reconstruction analysis relied on the presence of vGluT1 and vGluT2 immunostaining, we cannot exclude the possibility that GluD1 ablation alters the expression of these vesicular transporters, potentially resulting in synapses that were not labeled and therefore not included in our reconstructions. Additional presynaptic mechanisms not examined in this study should also be considered as potential contributors to GluD1 KO-mediated changes in thalamostriatal synaptic strength. For instance, changes in the number, size and localization of synaptic vesicles (Colliver et al., 2000; Edwards, 2007; Gong et al., 2003; Pothos, 2002; Sulzer & Edwards, 2000; Zhang et al., 1998) and mitochondrial energy supply (Safiulina et al., 2006; Verstreken et al., 2005; Villalba & Smith, 2010) are two important regulators of synaptic physiology that have been identified in various brain regions. Additionally, postsynaptic mechanisms besides the volume of spines and PSD areas should be considered. The morphology of striatal MSNs in KO mice has not yet been examined. A reduction in dendritic spines density as well as the decreased complexity and length of the dendritic arbor of individual MSNs could provide an alternative explanation for decreased excitatory input, although it remains unclear whether such changes would arise form presynaptic terminal loss or intrinsic alterations in postsynaptic structure.

#### Technical Considerations

It is noteworthy that our ultrastructural data were collected from a subset of striatal glutamatergic terminals at a single time point, and thus the functional and/or physiological implications should be interpreted with caution. Furthermore, although a substantial number of terminals and their synapses were reconstructed in 3D, the number of animals per group was relatively small (N = 3). Some variability was observed across and within individual animals. With a small size and non-normal distribution, nonparametric statistical analysis was used, which may limit the statistical power to detect subtle differences in synaptic ultrastructure between WT and KO mice. Despite these limitations, the data presented in Supplementary Figure 1 and 2 consistently support the conclusion that morphometric changes in thalamostriatal terminals and their postsynaptic targets are unlikely to be a primary driver of reduced thalamic excitatory input to MSNs.

## 5 CONCLUDING REMARKS

Despite growing interest in GluD1 and its relevance to neuropsychiatric disorders, its role in the mammalian CNS remains poorly understood. To further define GluD1 function in the striatum, our 3D reconstruction analysis revealed no significant ultrastructural differences that could serve as a morphological basis for the previously reported physiological reduction in thalamic input in mice with striatal-specific GluD1 ablation (Liu et al., 2020). These findings suggest that the physiological deficit is likely due to a loss of Pf terminals rather than changes in the ultrastructure of individual thalamostriatal synapses. Together, these results support a model in which GluD1 regulates input-specific circuit organization and synaptic connectivity, rather than the structural morphology of individual synapses. This interpretation is consistent with the emerging view that GluD1 functions as a trans-synaptic organizer within activity-dependent signaling complexes that regulate synapse formation and circuit function. Disruption of this mechanism may lead to circuit-level dysfunction without overt ultrastructural abnormalities, providing a framework for understanding behavioral inflexibility—a core symptom of autism and schizophrenia—in relation to GluD1 ablation in the dorsolateral striatum (Liu et al., 2020) and lesions or deactivation of the Pf-striatal system (Bradfield & Balleine, 2017). Future studies examining MSN morphology, including dendritic architecture and spine density, will be important for further defining the role of GluD1 in striatal circuitry.

