## Supplemental Figure 1 for "Effects of Glutamate Delta 1 Receptor (GluD1) Deletion on the Ultrastructural Features of Corticostriatal and Thalamostriatal Synapses in Mice"

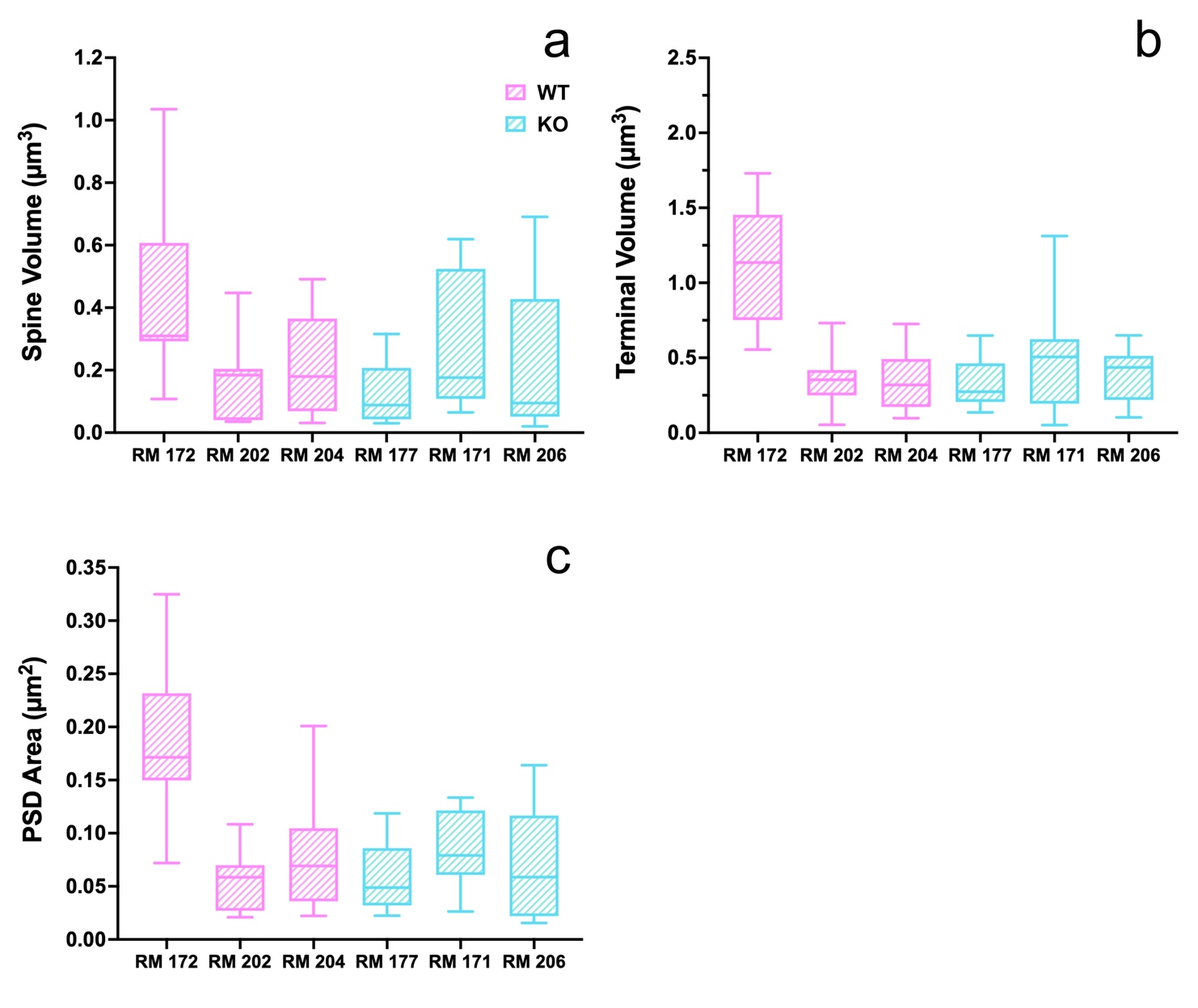

**Supplementary Figure 1:** Comparative analysis of axo-spinous synapses formed by vGluT1-IR terminals across individual animals in WT (red bars) and KO (blue bars) mice. (a–c) Box plots of dendritic spine volume (a), terminal volume (b), and PSD area (c) of axo-spinous synapses across 3 WT and 3 KO mice used in 3D reconstruction analysis. Data are presented separately for each animal to illustrate within-group variability.
