## Supplemental Figure 2 for "Effects of Glutamate Delta 1 Receptor (GluD1) Deletion on the Ultrastructural Features of Corticostriatal and Thalamostriatal Synapses in Mice"

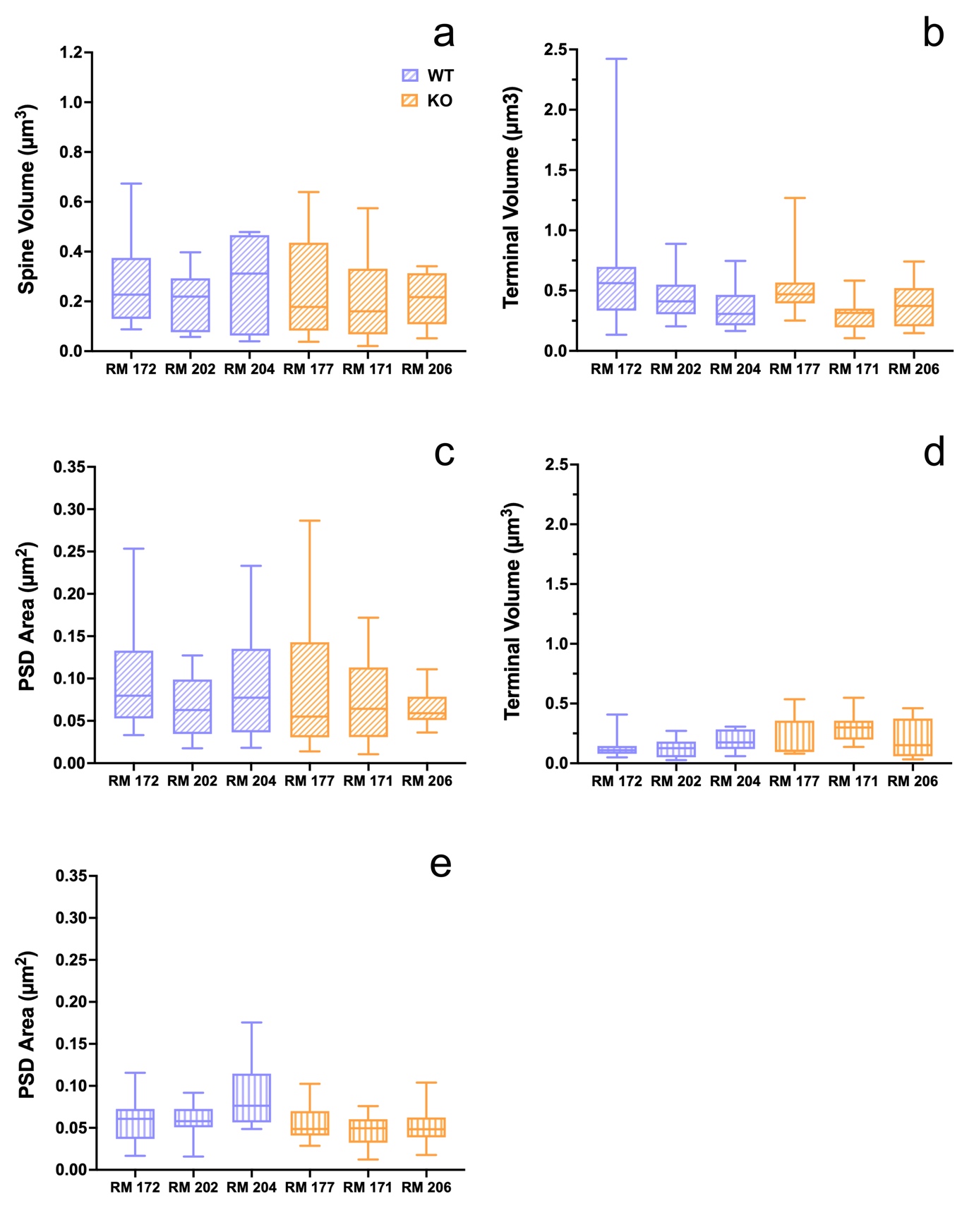

**Supplementary Figure 2:** Comparative quantitative analysis of axo-spinous and axo-dendritic synapses formed by vGluT2-IR terminals across individual animals in WT (purple bars) and KO (orange bars) mice. (a–c) Box plots of dendritic spine volume (a), terminal volume (b), and PSD area (c) of axo-spinous synapses across 3 WT and 3 KO mice used in 3D reconstruction analysis. (d and e) Box plots of terminal volume (d) and PSD area (e) of axo-dendritic synapses across 3 WT and 3 KO mice used in 3D reconstruction analysis. Data are presented separately for each animal to illustrate within-group variability.
