## Supplemental Figure 3 for "Effects of Glutamate Delta 1 Receptor (GluD1) Deletion on the Ultrastructural Features of Corticostriatal and Thalamostriatal Synapses in Mice"

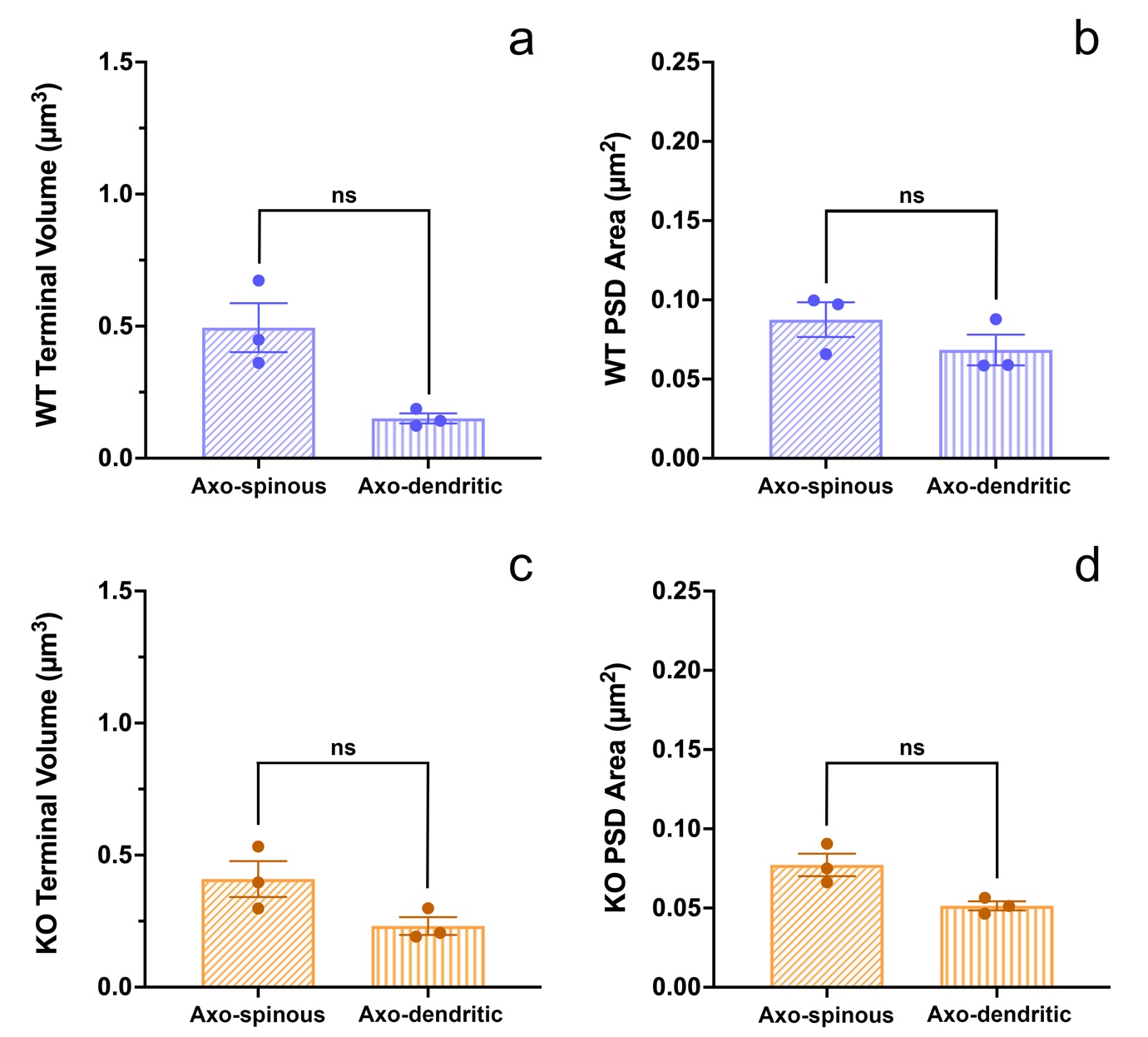

**Supplementary Figure 3:** Comparative quantitative analysis of terminal volume and PSD area in synapses formed by vGluT2-IR terminals. (a) In WT mice, the volume of terminals forming axo-spinous synapses is larger than those involved in axo-dendritic synapses, though this difference is not statistically significant (Wilcoxon signed-rank test, p = 0.25). (b) In WT mice, PSD areas of axo-spinous and axo-dendritic synapses are not significantly different (Wilcoxon signed-rank test, p = 0.25). (c) In KO mice, the volume of terminals forming axo-spinous synapses is slightly larger than those forming axo-dendritic synapses, though this difference is not statistically significant (Wilcoxon signed-rank test, p = 0.50). (d) In KO mice, no significant difference was found in PSD area between axo-spinous and axo-dendritic synapses (Wilcoxon signed-rank test, p = 0.25).
